# The U1 snRNP subunit LUC7 controls plant development and stress response through alternative splicing regulation

**DOI:** 10.1101/150805

**Authors:** Marcella de Francisco Amorim, Eva-Maria Willing, Anchilie G. Francisco-Mangilet, Irina Droste-Borel, Boris Maček, Korbinian Schneeberger, Sascha Laubinger

## Abstract

Introns are removed by the spliceosome, a large macromolecular complex composed of five ribonucleoprotein subcomplexes (U snRNP). The U1 snRNP, which binds to 5’ splice sites, plays an essential role in early steps of the splicing reaction. Here, we show that Arabidopsis LUC7 proteins, which are encoded by a three-member gene family in Arabidopsis, are important for plant development and stress resistance. We show that LUC7 are U1 snRNP accessory proteins by RNA immunoprecipitation experiments and LUC7 protein complex purifications. Transcriptome analyses revealed that LUC7 proteins are not only important for constitutive splicing, but also affects hundreds of alternative splicing events. Interestingly, LUC7 proteins specifically promote splicing of a subset of terminal introns. Splicing of LUC7-dependent introns is a prerequisite for nuclear export and some splicing events are modulated by stress in a LUC7-dependent manner. Taken together our results highlight the importance of the U1 snRNP component LUC7 in splicing regulation and suggest a previously unrecognized role of a U1 snRNP accessory factor in terminal intron splicing.

## Introduction

Eukaryotic genes are often interrupted by non-coding sequences called introns that are removed from pre-mRNAs while the remaining sequence, the exons, are joined together. This process, called splicing, is an essential step before the translation of the mature mRNAs and it offers a wide range of advantages for eukaryotic organisms. For instance, alternative splicing allows the production of more than one isoform from a single gene expanding the genome coding capacity (Kornblihtt et al., 2013; Reddy et al., 2013). Alternative splicing can also regulate gene expression by generating transcripts with premature termination codons (PTC) or/and a long 3’UTR, which may lead to RNA degradation via the nonsense-mediated decay (NMD) pathway (Kalyna et al., 2012; Drechsel et al., 2013; Shaul, 2015). Furthermore, splicing is usually coupled with other RNA processing events, such as 3’end formation and RNA transport to the cytosol (Kaida, 2016; Muller-McNicoll et al., 2016). In plants, alternative splicing contributes to essentially all aspects of development and stress responses (Carvalho et al., 2013; Staiger and Brown, 2013).

Intron removal is catalyzed by a large macromolecular complex, the spliceosome, which is formed by five small ribonucleoprotein particles (U snRNP): the U1, U2, U4, U5 and U6 snRNP. Each U snRNP contains a heteroheptameric ring of Sm or Lsm proteins, snRNP-specific proteins and an uridine-rich snRNA. Additional non-core spliceosomal proteins participate during the splicing reaction affecting exon-intron recognition and thus splicing efficiency. The canonical splicing cycle starts with binding of the U1 snRNP to the 5’ splice site (5’ss), followed by association of auxiliary proteins such as U2AF to the pre-mRNA, which facilitate the recognition of the 3’ splice site (3’ss). The thereby formed complex E recruits the U2 snRNP to generate complex A. In the next step, a trimeric complex consisting of U4/U5/U6 snRNPs joins to form complex B. Several rearrangements and ejection of the U1 and U4 snRNP are necessary to generate a catalytically active splicing complex (Wahl et al., 2009; Will and Luhrmann, 2011).

The fact that U1 snRNP is recruited to the 5’ss in the initial step of splicing suggests that this complex is necessary for the correct 5’ splicing site selection. Indeed, it has been shown that U1-deficient zebrafish mutants accumulate alternative spliced transcripts, suggesting that the U1 snRNP indeed fulfills regulatory roles in splice site selection (Rosel et al., 2011). Although the spliceosome consists of stoichiometrically equal amounts of each subunit, the U1 snRNP is more abundant than all the other spliceosomal subcomplexes (Kaida et al., 2010; Kaida, 2016). One reason for this is that the U1 snRNP executes splicing independent functions. The metazoan U1 snRNP, for instance, binds not only to the 5’ss, but also throughout the nascent transcript blocking a premature cleavage and polyadenylation (Kaida et al., 2010; Berg et al., 2012). Furthermore, the U1 snRNP is important to regulate promoter directionality and transcription in animals (Almada et al., 2013; Guiro and O’Reilly, 2015).

U1 snRNP complexes were purified and characterized in yeast and human. The U1 snRNP contains the U1 snRNA, Sm proteins, three U1 core proteins (U1-70K, U1-A and U1-C) and U1-specific accessory proteins, such as LUC7, PRP39 and PRP40. All these proteins are conserved in plants suggesting a U1 snRNP composition very similar to the one in yeast and metazoans (Wang and Brendel, 2004; Koncz et al., 2012; Reddy et al., 2013). Interaction studies revealed that U1 snRNP associates with serine-arginine (SR) proteins, indicating a complex mechanism for splicing site selection that involves also non-snRNP proteins (Golovkin and Reddy, 1998; Cho et al., 2011).

The function of the plant U1 snRNP is not well characterized. This might be due to the fact that in *Arabidopsis thaliana* the core U1 snRNP components *U1-70K* or *U1C* are single copy genes and a complete knockout most likely causes severe mutant phenotypes or lethality. On the other hand, proteins such as PRP39, PRP40 and LUC7 are encoded by small gene families, which require the generation of multiple mutants for functional studies. Some U1 specific Arabidopsis mutants have been characterized: Mutations in the accessory factor PRP39A cause delayed flowering due to increased expression of the flowering time regulator *FLOWERING LOCUS C* (*FLC*), but the mutants do not exhibit severe developmental defects (Wang et al., 2007; Kanno et al., 2017). In a reverse genetic approach, *U1-70K* expression was specifically reduced in flowers by an antisense RNA and the resulting transgenic plants exhibit strong floral defects (Golovkin and Reddy, 2003). Moreover, a mutation in U1A causes an altered salt stress response (Gu et al., 2017). Thus, despite evidences that U1 snRNP is essential for plant development and stress response, the functions of the U1 snRNP in regulating the transcriptome of plants are largely unknown. Other characterized factors, such as GEMIN2 or SRD2 are required for the functionality of all snRNPs, but not specifically for U1 function (Ohtani and Sugiyama, 2005; Schlaen et al., 2015).

Here, we report on the functional characterization of *Arabidopsis* mutants impaired in U1 snRNP function. For this, we focused in this study on the U1 snRNP components LUC7, which we show to be essential for normal plant development and plant stress resistance. Our whole transcriptome analyses on *luc7* triple mutant show that impairments of LUC7 proteins affect constitutive and alternative splicing. Surprisingly, our results reveal the existence of transcripts, in which terminal introns are preferentially retained in a LUC7-dependent manner. Unspliced LUC7-dependent introns cause a nuclear retention of the pre-mRNAs and the splicing efficiency of LUC7-dependent introns can be modulated by stress. Our results suggests that the plant U1 snRNP components LUC7 regulate alternative splicing of pre-mRNAs and thereby impact their nuclear export, which could be a mechanism to fine-tune gene expression under stress conditions.

## Results

### LUC7 proteins, a family of conserved nuclear zinc-finger / arginine-serine (RS) proteins, redundantly control plant development

*LETHAL UNLESS CBC 7* (*LUC7*) was first identified in a screen for synthetic lethality in a yeast strain lacking the nuclear cap-binding complex (CBC), which is involved in several RNA processing events (Fortes et al., 1999a; Gonatopoulos-Pournatzis and Cowling, 2014; Sullivan and Howard, 2016). LUC7 proteins carry a C_3_H and a C_2_H_2_-type zinc-fingers, which are located in the conserved LUC7 domain. LUC7 proteins from higher eukaryotes usually contain also an additional C-terminal Arginine/Serine-rich (RS) domain, which is known to mediate protein-protein interactions (Puig et al., 2007; Webby et al., 2009; Heim et al., 2014). *Arabidopsis thaliana* encodes three *LUC7* genes (*AthLUC7A*, *AthLUC7B* and *AthLUC7RL*), which are separated in two clades: *LUC7A/B* and *LUC7RL* (Figure 1A and S1). AthLUC7RL is more similar to its yeast homolog and lacks a conserved stretch of 80 amino acids of unknown function present in AthLUC7A and AthLUC7B (Figure S1). A phylogenetic analysis revealed that algae contain a single *LUC7* gene belonging to the LUC7RL clade reinforcing the idea that LUC7RL proteins are closer to the ancestral LUC7 than LUC7A/B. In the moss Physcomitrella and in the fern Selaginella one can find proteins belonging to both clades, suggesting that the split into *LUC7RL* and *LUC7A/B* occurred early during the evolution of land plants.

**Figure 1:**
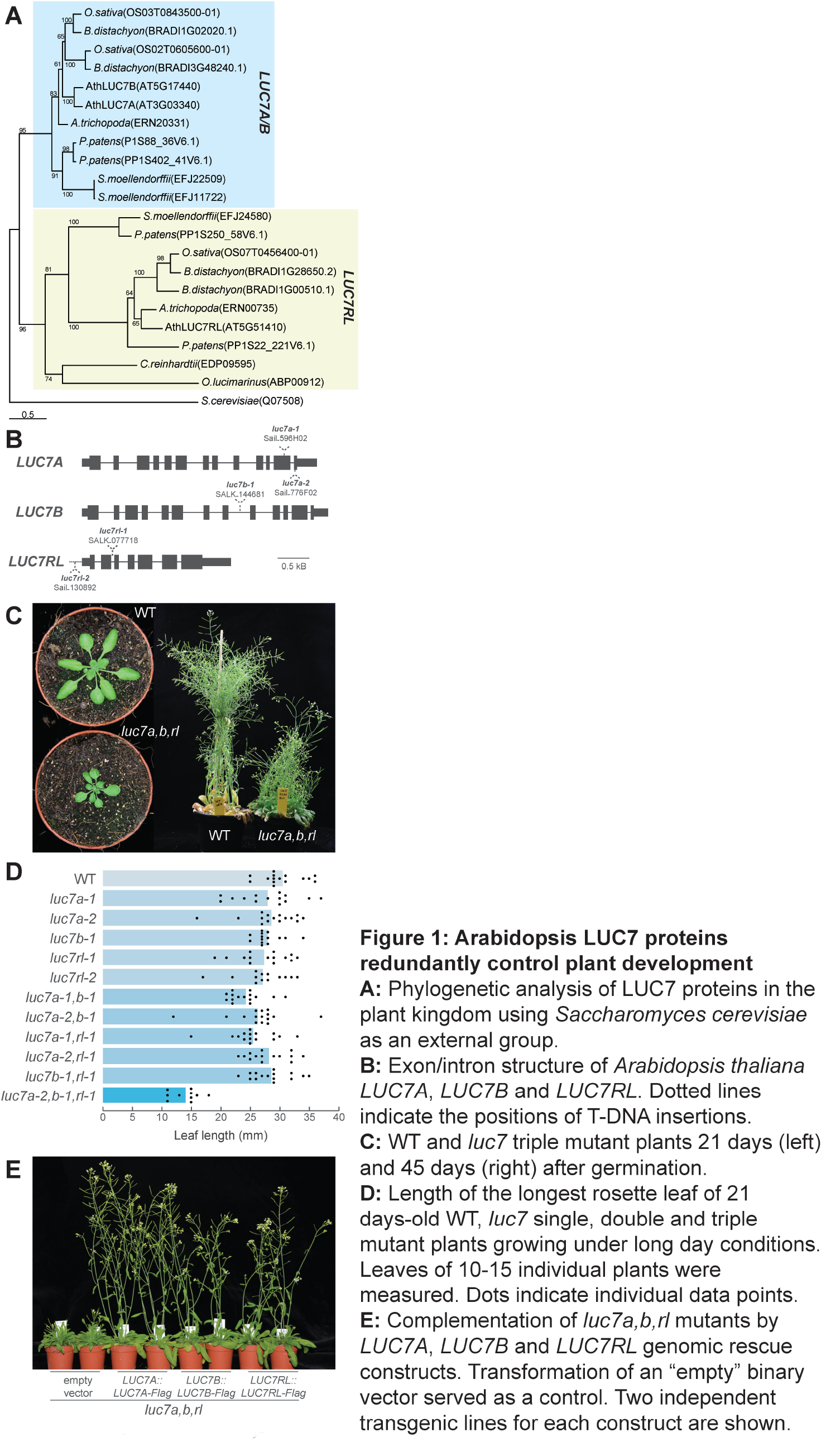
*Arabidopsis* LUC7 proteins redundantly control plant development. **A:** Phylogenetic analysis of LUC7 proteins in the plant kingdom using *Saccharomyces cerevisiae* as an external group. **B:** Exon/intron structure of *Arabidopsis thaliana LUC7A*, *LUC7B* and *LUC7RL*. Dotted lines indicate the positions of T-DNA insertions. **C:** WT and *luc7* triple mutant plants 21 days (left) and 45 days (right) after germination. **D:** Length of the longest rosette leaf of 21 days-old WT, *luc7* single, double and triple mutant plants growing under long day conditions. Leaves of 10-15 individual plants were measured. Dots indicate individual data points. **E:** Complementation of *luc7a,b,rl* mutants by *LUC7A*, *LUC7B* and *LUC7RL* genomic rescue constructs. Transformation of an “empty” binary vector served as a control. Two independent transgenic lines for each construct are shown.

In order to understand the function of the Arabidopsis U1 snRNP, we analyzed T-DNA insertion lines affecting *LUC7* genes (Figure 1B). Single and double *luc7* mutants were indistinguishable from wild-type plants (WT) (Figure S2). However, *luc7* triple mutant exhibit a wide range of developmental defects, including dwarfism and reduced apical dominance (Figure 1C-E). To test whether the impairment of *LUC7* functions was indeed responsible for the observed phenotypes, we reintroduced a wild-type copy of *LUC7A*, *LUC7B* or *LUC7RL* in the *luc7* triple mutant. Each of the *LUC7* genes was sufficient to restore the growth phenotype of the *luc7* triple mutant (Figure 1E). These results reveal that the phenotype observed in this mutant is attributable to the impairment of *LUC7* function and it suggests that *LUC7* genes act redundantly to control *Arabidopsis* growth and development.

### LUC7 functions in the ABA pathway and is important for cold and salt stress responses

Splicing is essential for plant stress resistance and mutants impaired in splicing often react differently to stress and the stress hormone abscisic acid (ABA) (Filichkin et al., 2015; Zhan et al., 2015). In addition, global impairment of the splicing machinery elicits ABA signaling (AlShareef et al., 2017; Ling et al., 2017). To test whether LUC7 is important for plant stress resistance and ABA-mediated stress signaling, we analyzed growth parameters of WT, the *luc7* triple mutant and a *luc7* rescue line in presence of exogenous ABA or salt. A cotyledon greening assay showed that *luc7* triple mutants reacted hypersensitively to exogenous ABA (Figure 2A, B), suggesting that *LUC7* plays an important role in the ABA pathway. Furthermore, salt in the growth medium impaired root growth much more strongly in *luc7* triple mutant than in WT or in a *luc7* rescue line (Figure 2C, D). Similarly, cold temperatures strongly compromised the growth of *luc7* triple mutants when compared to WT (Figure 2E). These results imply that functional *LUC7* proteins are required for plant stress resistance and ABA responses.

**Figure 2:**
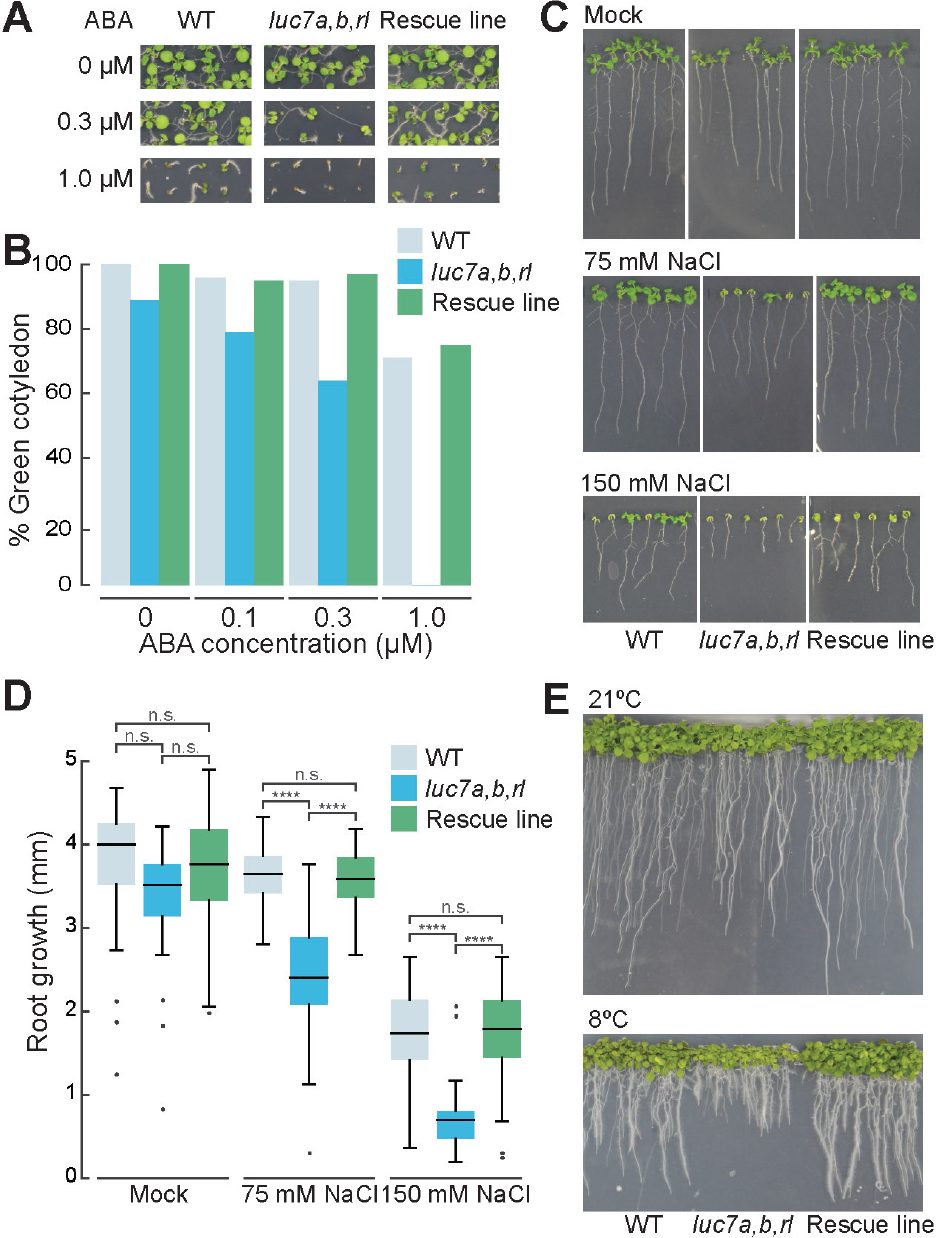
*Arabidopsis* LUC7 is involved in ABA signaling and salt stress responses **A,B:** WT, *luc7* triple mutant and a *luc7* rescue line (*luc7a,b,rl; pLUC7A:LUC7A-YFP*) were grown on half-strength MS plates containing 1% sucrose and indicated amount of ABA. Seedling phenotypes (A) and quantification of seedlings with green cotyledons (B) are shown. Green cotyledons were scored ten days after germination. One of two biological replicates is shown. **C,D:** WT, *luc7* triple mutant and a *luc7* rescue line (*luc7a,b,rl; pLUC7A:LUC7A-YFP*) were germinated on half-strength MS vertical plates and seedling were transferred on half-strength MS plates containing the indicated amount of NaCl. Plates were always placed vertically and the root growth was scored over 7 days. Phenotypes (C) and root length quantification (D) are shown. **E:** Gross phenotype of WT, *luc7* triple mutant and a *luc7* rescue line grown at 22°C and 8°C.

### LUC7 is a U1 snRNP component in plants

The composition of the U1 snRNP subcomplex is known in yeast and metazoans but not in plants (Will and Luhrmann, 2001; Koncz et al., 2012). Therefore, we asked whether LUC7 is also an U1 component in Arabidopsis. Due to the fact that our genetic analyses of *luc7* mutants suggested that LUC7 proteins act largely redundant, we focused our further analyses mainly on a single LUC7 protein, LUC7A. A protein that is part of the U1 complex is tightly associated with U1 specific components such as the U1 snRNA. To test whether LUC7 is found in a complex with the U1 snRNA, we performed RNA immunoprecipitation (RIP) experiments using a *luc7* triple mutant carrying *pLUC7A:LUC7A-YFP* rescue construct (Figure S3). Immunoprecipitation of LUC7A-YFP enriched the U1 snRNA more than 40-fold, but did not enrich two unrelated, but abundant RNAs, U3 snoRNA and *ACTIN* mRNA (Figure 3A). Small amounts of U2 snRNA was also found associated with LUC7A, which is in agreement with the fact that U1 and U2 snRNP directly interact to form spliceosomal complex A (Figure 3A). However, the amount of recovered U2 snRNA is more than four-fold lower than that of the U1 snRNA (Figure 3A). These results strongly suggest that Arabidopsis LUC7 proteins are bona fide U1 snRNP components.

**Figure 3:**
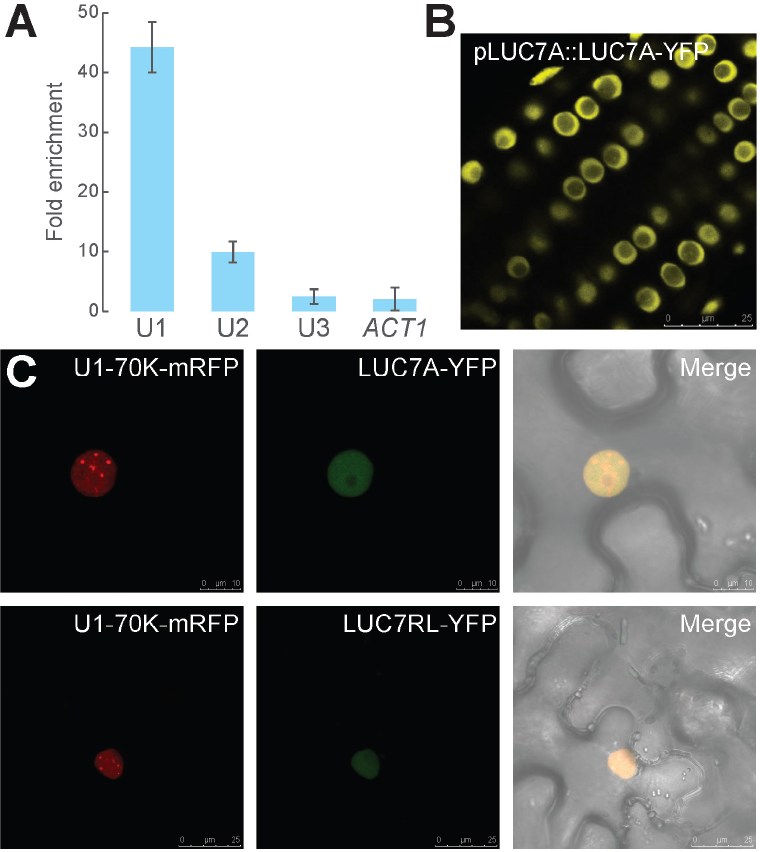
*Arabidopsis* LUC7 is an U1 snRNP component **A:** RNA immunoprecipitation using a *pLUC7A:LUC7A-YFP, luc7a,b,rl* complemented line. Proteins were immunoprecipitated using GFP-specific affinity matrix and RNAs were extracted from the input and the immunoprecipitated fraction. U1, U2, U3 snRNAs and *ACTIN* RNA were quantified using qRT-PCR. Enrichment of the respective RNA in *LUC7A:LUC7A-YFP luc7a,b,rl* transgenic line was calculated over WT (negative control). Error bars denote the range of two biological replicates. **B:** Subcellular localization of LUC7A in *pLUC7A:LUC7A-YFP luc7a,b,rl* in Arabidopsis transgenic plants. Roots of 9 day-old seedlings were analyzed using confocal microscopy. Scale bar indicates 25 µm. **C:** U1-70K-mRFP and LUC7A-YFP or LUC7RL-YFP proteins were transiently expressed in *N. benthamiana*. The subcellular localization of mRFP and YFP fusion proteins was analyzed using confocal microscopy. Scale bars indicate 10 µm and 25 µm for upper and lower panel, respectively.

Next we analyzed the subcellular localization of LUC7A and its co-localization with a core U1 snRNP subunit. LUC7A localized to the nucleus, but not to the nucleolus in *Arabidopsis* plants containing the *pLUC7A:LUC7A-YFP* rescue construct (Figure 3B). In addition, LUC7A partially co-localized with U1-70K in the nucleoplasm when transiently expressed in *Nicotiana benthamiana* (Figure 3C). Similar results were obtained for LUC7RL, the Arabidopsis LUC7 most distant in sequence to LUC7A (Figure 3C). In plants, co-localizations studies in protoplasts have shown that also the core U1 components only partially colocalize (Lorkovic and Barta, 2008). These partial colocalizations suggest that plant U1 snRNP proteins may fulfill additional functions as it has been observed in other eukaryotes (Workman et al., 2014).

To further test whether LUC7A associates in planta with known U1 snRNP components, we purified LUC7A-containing complexes. For this, we used *pLUC7A:LUC7A-YFP* complemented lines and as controls wild-type plants and transgenic lines expressing free GFP (*p35S:GFP*). Immunopurifications (IPs) were carried out three to four time independently. We observed that WT often produced more background in mass spectrometry (MS) analyses than the 35S:GFP line and we therefore decided to use WT as a more stringent control (Table S1). Among all identified proteins we considered those putative LUC7 interactors that were found in at least two independent experiment and were at least three time more abundant in *pLUC7A:LUC7A-YFP* IPs than in WT IPs. The mass spectrometry (MS) analysis revealed that LUC7 is indeed found in a complex with core U1 snRNP proteins U1A and U1-70K (Table 1, Table S1). Moreover, we detected peptides corresponding to the spliceosomal complex E components U2AF35 and U2AF65, further suggesting that LUC7 proteins are involved in very early steps of the splicing cycle (Table 1, Table S1). Additional proteins known to be involved in splicing and general RNA metabolism including several serine-arginine (SR) proteins (SR30, SCL30A, SCL33), SR45, SERRATE (SE) and the CBC component ABH1/CBP80 were found in LUC7A-containing complexes (Table 1, Table S1). To test the validity of the LUC7 complex purification experiment, we confirmed the interaction between LUC7 with SE and ABH1/CBP80 by in planta co-immunoprecipitation experiment (Figure S4). Interestingly, we also identified regulatory proteins in LUC7A-containing complexes, among them several kinases and proteins involved in 3’end processing (Table 1, Table S1).

**Table 1:**
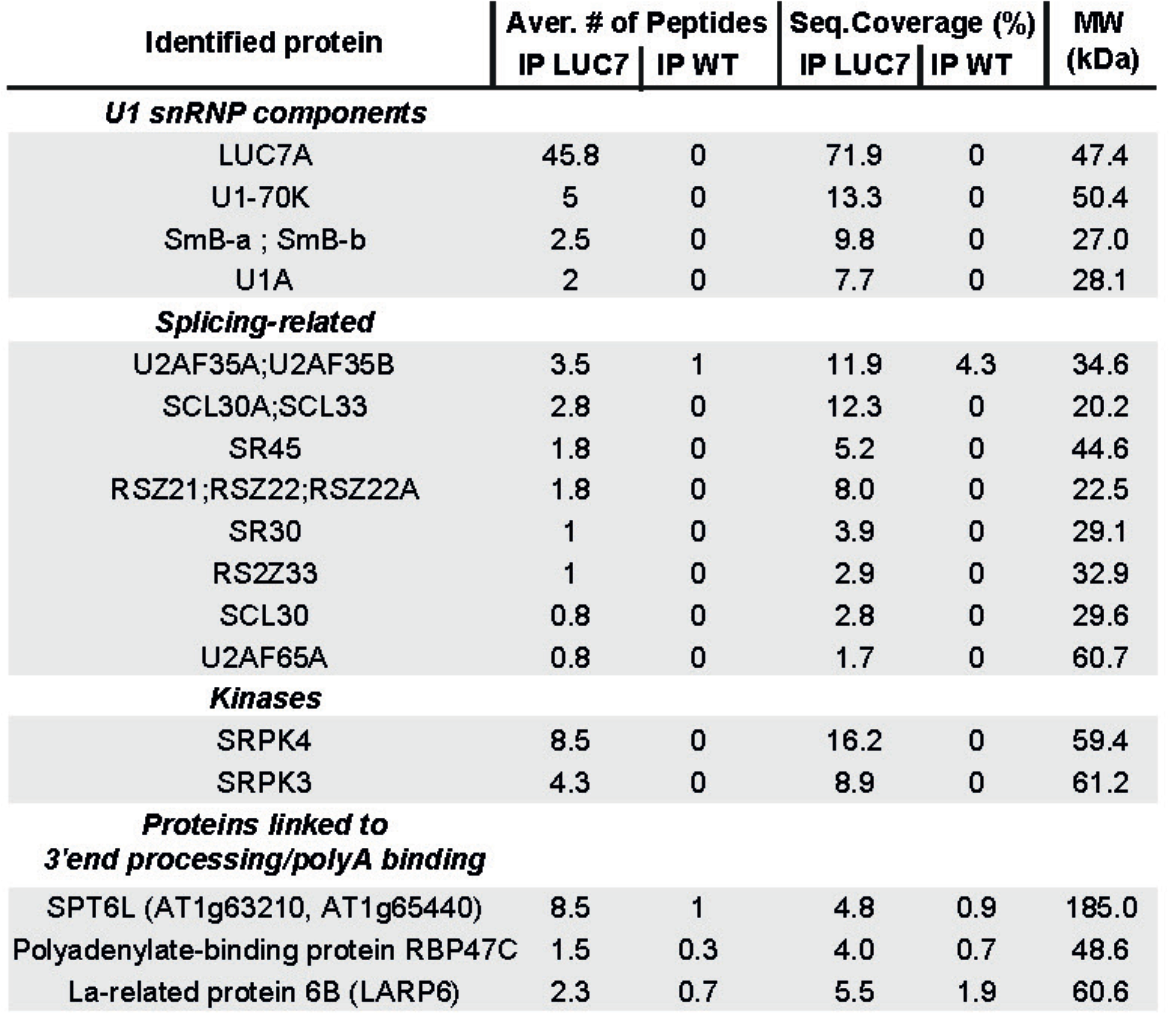
List of selected potential LUC7A interacting proteins identified in immunoprecipitation experiments followed by MS analysis.

### *LUC7* effects on the Arabidopsis coding and non-coding transcriptome

In order to identify misregulated and misspliced genes in *luc7* mutants, we performed an RNA-sequencing (RNA-seq) analysis with three biological replicates. We decided to use seven days old WT and *luc7* triple mutant seedlings. At this age, *luc7* triple mutant and WT seedlings are morphologically similar and therefore, changes in transcript levels and splicing patterns most likely reflect changes caused by impairments of LUC7 proteins and are not due to different contribution of tissues caused by, for instance, a delay in development or/and different morphology (Figure S5). We sequenced between 22.1 and 27.6 million reads per library.

An analysis of differentially expressed genes revealed that 840 genes are up- and 703 are downregulated in *luc7* triple mutant when compared to WT (Table S2, Table S3). The majority of genes that change expression were protein-coding genes (Figure 4A). Nevertheless, non-coding RNAs (ncRNAs) were significantly enriched among the *LUC7* affected genes (p < 0.05, hypergeometric test), although the overall number of ncRNA affected in *luc7* triple mutant is relatively small (Figure 4A, B). Previous studies implied that the U1 snRNP regulates microRNA (miRNA) biogenesis (Bielewicz et al., 2013; Schwab et al., 2013; Knop et al., 2016; Stepien et al., 2017). However, the expression of *MIRNA* genes was not affected in *luc7* triple mutants (Figure 4A). In addition, quantification of mature miRNA levels revealed that all tested miRNAs did not change abundance in *luc7* triple mutants (Figure 4C). These results show that LUC7 proteins affect the expression of protein-coding genes and a subset of ncRNAs, but are not involved in the miRNA pathway.

**Figure 4:**
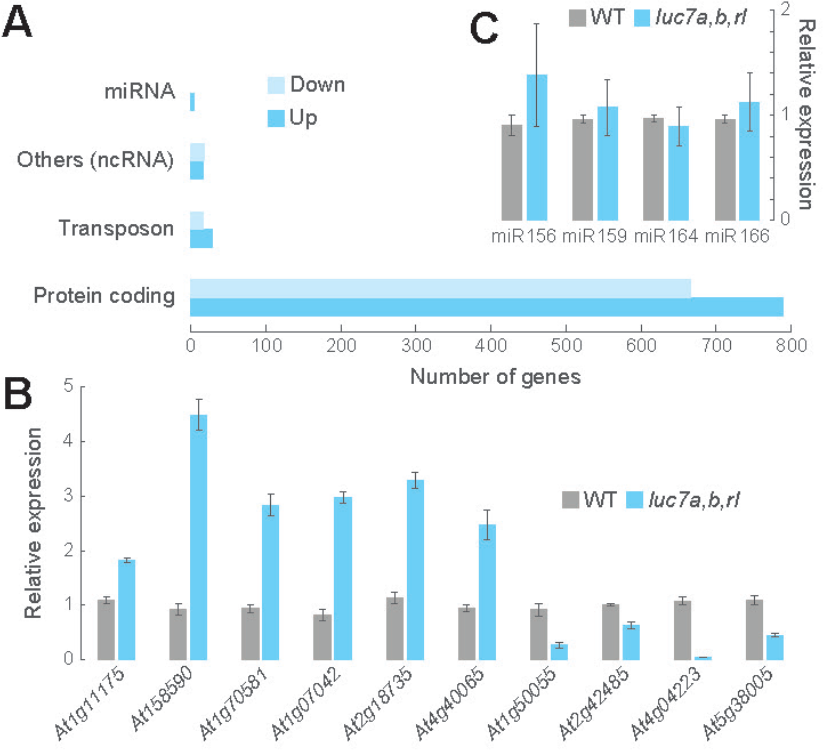
Mutations in *LUC7* result in misexpression of protein-coding and non-coding genes, but not in *MIRNA* genes. **A:** Differentially expressed genes in *luc7a,b,rl* mutant compared to WT. **B,C:** qRT-PCR analysis of selected ncRNA **(B)** and miRNAs **(C)** in WT and *luc7a,b,rl*. Error bars denote the SEM (n=3).

### Arabidopsis *LUC7* functions are important for constitutive and alternative splicing

Because LUC7 proteins are U1 snRNP components, we ask whether misspliced transcripts accumulate in the *luc7* triple mutant. In total, we identified 640 differential splicing events in *luc7* triple mutants compared to WT (Table S4). Only 17 of these alternative splicing events occurred in mRNAs whose expression also differed between *luc7* mutants and WT (Table S5, Table S6). Hence, the splicing differences found were mainly not due to changes in transcript abundance. We detected a large number of intron retention events in the *luc7* triple mutant (Figure 5A). RT-PCR experiments with oligonucleotides flanking selected intron retentions events confirmed the RNA-seq data (Figure 5B). These results suggest that lack and/or impairment of the U1 snRNP component LUC7 disturbs intron recognition and thus splicing. We also identified a large number of exons skipping events in the *luc7* triple mutant. Exon skipping is also a major outcome of impairing U1 snRNP function or binding in metazoans (Lorkovic and Barta, 2008; Rosel et al., 2011). These defects are most likely caused by interactions of the U2 snRNP with U1 snRNPs associated to alternative 5’splice sites (Morcos, 2007). Furthermore, we detected several cases of alternative 5’ and 3’ splice site selection in *luc7* triple mutants (Figure 5A-F).

**Figure 5:**
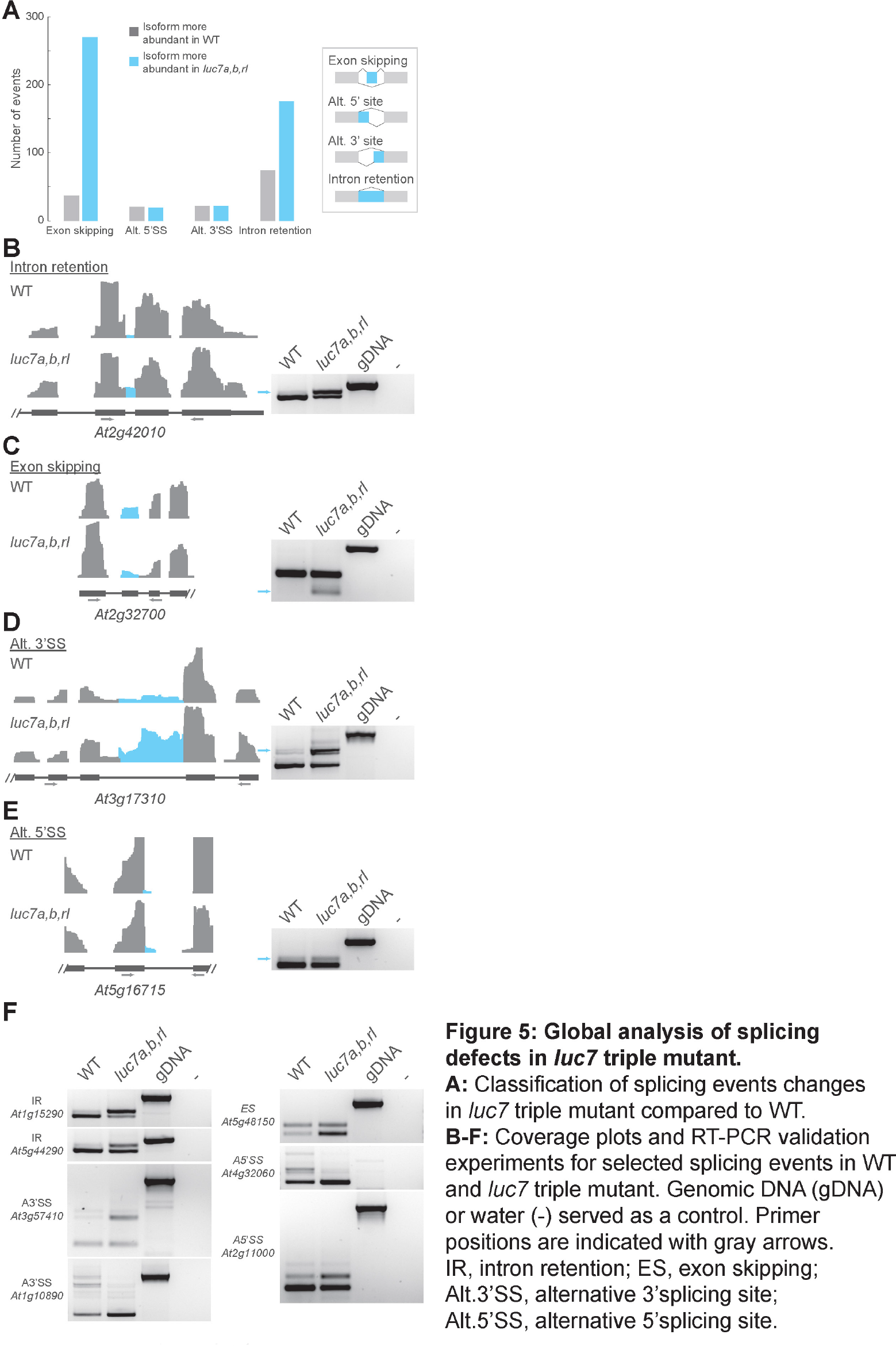
Global analysis of splicing defects in *luc7* triple mutant. **A:** Classification of splicing events changes in *luc7* triple mutant compared to WT. **B-F:** Coverage plots and RT-PCR validation experiments for selected splicing events in WT and *luc7* triple mutant. Genomic DNA (gDNA) or water (-) served as a control. Primer positions are indicated with gray arrows. IR, intron retention; ES, exon skipping; Alt.3’SS, alternative 3’splicing site; Alt.5’SS, alternative 5’splicing site.

Some splicing defects observed in *luc7* triple mutants generated transcript variants that did not exist in WT (e.g. *At2g32700*, Figure 5C). In these cases, LUC7 proteins affect the splicing of an intron which is constitutively removed in WT plants. On the contrary, in other cases the *luc7* triple mutant lacked specific mRNA isoforms, which exist in wild-type plants (e.g. *At1g10980*, *At4g32060*), or the ratio of two different isoforms was altered in *luc7* triple mutant when compared to WT (e.g. *At3g17310*, *At5g16715*, *At5g48150*, *At2g11000*) (Figure 5D-F). In these cases, LUC7 proteins affect a splicing event which is subjected to alternative splicing in WT plants. These results show that LUC7 proteins are involved in both constitutive and alternative splicing in Arabidopsis.

Next, we checked whether splicing changes observed in *luc7* triple mutant are actually due to the loss of only a specific *LUC7* gene or whether *LUC7* genes act redundantly. To test this, we analyzed the splicing pattern of some mRNAs in *luc7* single, double and triple mutants. Some splicing defects were detectable even in *luc7* single mutants (Figure S6), but the degree of missplicing increased in *luc7* double and triple mutants suggesting that LUC7 proteins act additively on these introns (e.g. *At5g16715*). Some splicing defects occurred only in *luc7* triple mutants, implying that LUC7 proteins act redundantly to ensure splicing of these introns (e.g. *At1g60995*). Other splicing defects might more likely be due to the lack of LUC7A/B or LUC7RL. For instance, intron removal of *At2g42010* more strongly relied on *LUC7RL*, while removal of an intron in *At5g41220* preferentially depends on LUC7A/LUC7B (Figure S6). These findings suggest that Arabidopsis *LUC7* genes function redundantly, additively or specifically to ensure proper splicing.

### LUC7 proteins are preferentially involved in the removal of terminal introns

In yeast, LUC7 connects the CBC with the U1 snRNP and this interaction is important for the correct 5’ splicing site selection (Fortes et al., 1999b). In plants, the CBC associates with SE to ensure splicing of cap-proximal first introns (Laubinger et al., 2008; Raczynska et al., 2010; Raczynska et al., 2013). In addition, we show here that LUC7A form complexes with SE and ABH1/CBP80, one of the CBC competent (Table S1, Figure S4). To investigate the relationship between LUC7 and the CBC/SE in plants, we analyzed the splicing patterns of LUC7 dependent introns in *cbc* mutants (*cbp20* and *cbp80*) and *se-1* by RT-PCR. All tested introns retained in *luc7* triple mutant were correctly spliced in *cbc* and *se* mutants (Figure 6A). Conversely, first introns that were partially retained in *cbp20, cbp80* and *se-1* mutants were completely removed in the *luc7* triple mutant (Figure 6B). These observations suggest that the functions of LUC7 and CBC/SE in splicing of the selected introns do not overlap.

**Figure 6:**
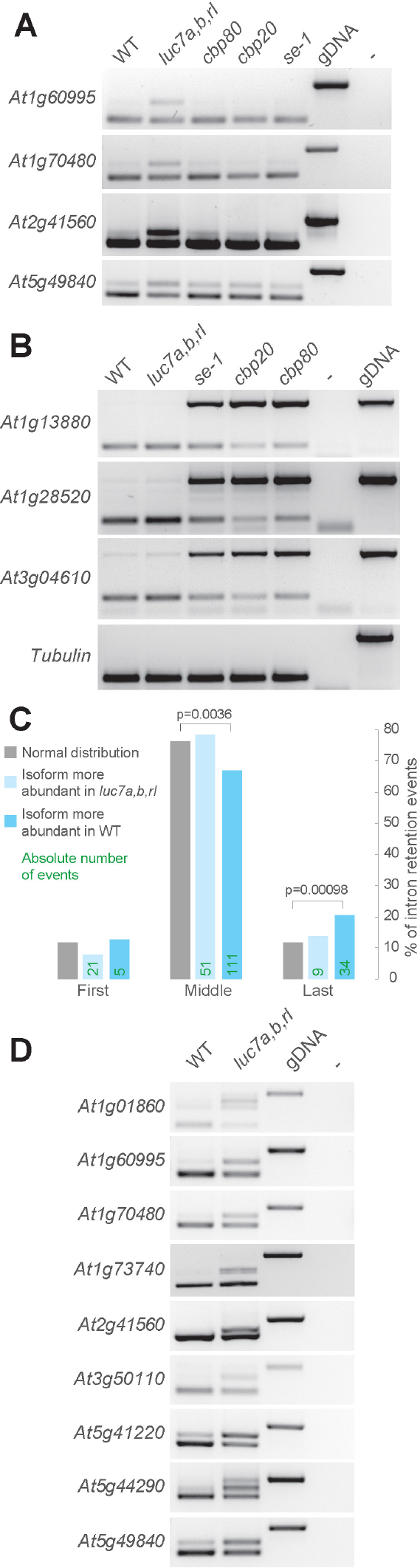
LUC7 proteins have a pronounced effect on terminal intron splicing. **A:** RT-PCR analysis of LUC7-dependent introns in WT, *luc7* triple mutant, *cbp80*, *cbp20* and *se-1* mutants. **B:** RT-PCR analysis of CBC/SE-dependent introns in WT, *luc7* triple mutant, *cbp80*, *cbp20* and *se-1* mutants. **C:** Classification of intron retention according to the intron position (first, middle, or last). Only genes with 3 or more introns were considered for this analysis. **D:** RT-PCR analysis of genes carrying retained terminal introns in WT and *luc7* triple mutants.

Next, we asked whether LUC7 has a preference for promoting splicing of cap-proximal first introns as it has the CBC/SE complex. We classified retained introns in *luc7* triple mutant according to their position within the gene (first, middle or last introns). Only genes with at least 3 introns were considered for this analysis. We found a significant increase in retained last introns, but not first introns, in *luc7* triple mutants (Figure 6C). Although the total number of retained introns was higher among middle introns, the relative amount of retained middle introns in *luc7* triple mutant was significantly reduced (Figure 6C). Retention of terminal introns in *luc7* triple mutants was confirmed by RT-PCR analysis (Figure 6D). In summary, our data revealed that (i) CBC/SE acts independently of LUC7 in splicing of cap-proximal introns and that (ii) LUC7 proteins play an important role for the removal of certain terminal introns.

### mRNAs harboring unspliced LUC7-dependent introns remain in the nucleus and escape NMD

We were further interested in determining the characteristics and possible functions of LUC7-dependent introns. When introns are retained, the resulting mRNA can contain a premature stop codon and a long 3’UTR, which are hallmarks of NMD targets (Kalyna et al., 2012; Drechsel et al., 2013; Shaul, 2015). To check whether mRNAs containing a retained LUC7-dependent introns are NMD substrates, we analyzed their splicing patterns in two mutants impaired in NMD, *lba-1* and *upf3-1*. If unspliced isoforms were indeed NMD targets, we would expect their abundance to be increased in NMD mutants. Interestingly, we did not observe any change between WT and *upf* mutants (Figure 7A). Thus, we conclude that the tested LUC7-dependent introns do not trigger degradation via the NMD pathway.

**Figure 7:**
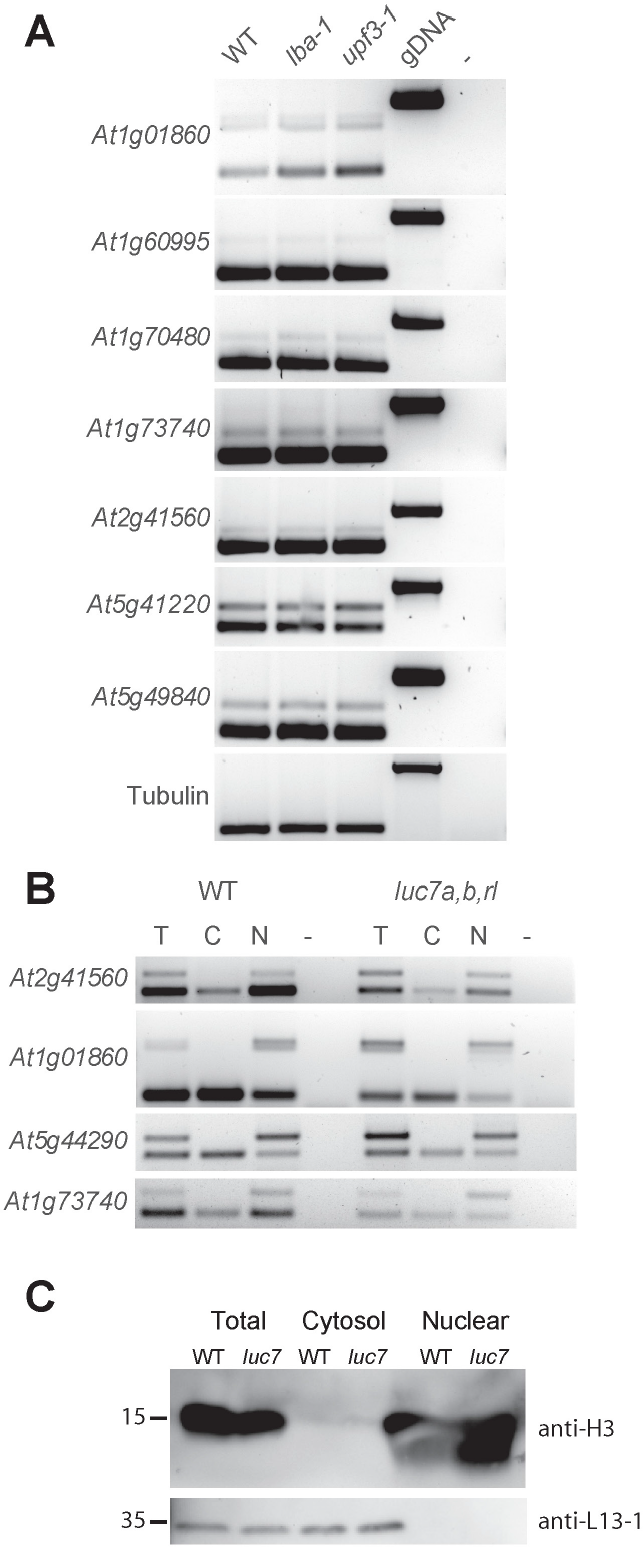
mRNAs containing retained LUC7-dependent introns are NMD-insensitive and remain nuclear. **A:** RT-PCR analysis of LUC7-dependent introns in WT and NMD mutants (*lba1* and *upf3-1*). **B:** Splicing patterns of mRNAs isolated from total (T), cytosolic (C) and nuclear (N) fractions. **C:** Immunoblot analysis of proteins isolated from total, cytosolic and nuclear fractions. Blots were probed with antibodies against histone H3 and a ribosomal protein, L13-1.

NMD occurs in the cytoplasm and RNAs can escape NMD by not being transported from the nucleus to the cytosol (Gohring et al., 2014). We therefore checked in which cellular compartment mRNAs with spliced and unspliced LUC7-dependent introns accumulate. To do this, we isolated total, nuclear and cytosolic fractions from wild-type and *luc7* triple mutant plants and performed RT-PCR analyses (Figure 7B). Purity of cytosolic and nuclear fractions was controlled by immunoblot analysis using antibodies against histone H3 (specific for nuclear fractions) and a 60S ribosomal protein (L13-1, specific for cytosolic fractions) (Figure 7C). Spliced mRNA isoforms accumulated in the cytosol, whereas mRNAs containing the unspliced LUC7-dependent introns were found in nuclear fractions (Figure 7B). These results indicate that retention of LUC7-dependent introns correlates with trapping mRNAs in the nucleus and suggest that splicing of LUC7-dependent introns is essential for mRNA transport to the cytosol.

### Splicing of LUC7-dependent introns can be modulated by stress

Our results revealed that a subset of alternatively spliced introns requires LUC7 proteins for efficient splicing and that splicing of these introns is a prerequisite for nuclear export. This mechanism could serve as a nuclear quality control step to prevent that unspliced mRNAs are exported prematurely. Interestingly, a GO analysis of genes containing LUC7-dependent introns indicated an enrichment for stress related genes (Figure S7). This prompted us to speculate that nuclear retention of mRNAs could be exploited as a regulatory mechanism to fine-tune gene expression under stress conditions.

To test this hypothesis, we decided to check the splicing of some LUC7-dependent introns in WT under stress condition. We chose cold stress because *luc7* mutants are cold-sensitive (Figure 2) and in addition, it was suggested that U1 snRNP functionality is impaired under cold condition (Schlaen et al., 2015). To quantify the amount of unspliced isoforms in cold condition, we designed qPCR-primers specific to unspliced isoforms and the total mRNA pool and calculate the relative amount of mRNA carrying unspliced LUC7-dependent introns compared to the total mRNA pool. mRNAs of *At1g70480*, *At2g41560* and *At5g44290* significantly accumulated unspliced isoforms in responses to cold treatment demonstrating that cold stress modulates the splicing efficiency of these LUC7-dependent introns (Figure 8A). Interestingly, the amount of unspliced mRNA in *luc7* triple mutants does not differ significantly between mock and stress conditions (Figure 8A). This observation suggests that LUC7 is directly involved in the regulation of intron splicing under stress conditions and that LUC7 might be a target for stress response pathways.

**Figure 8:**
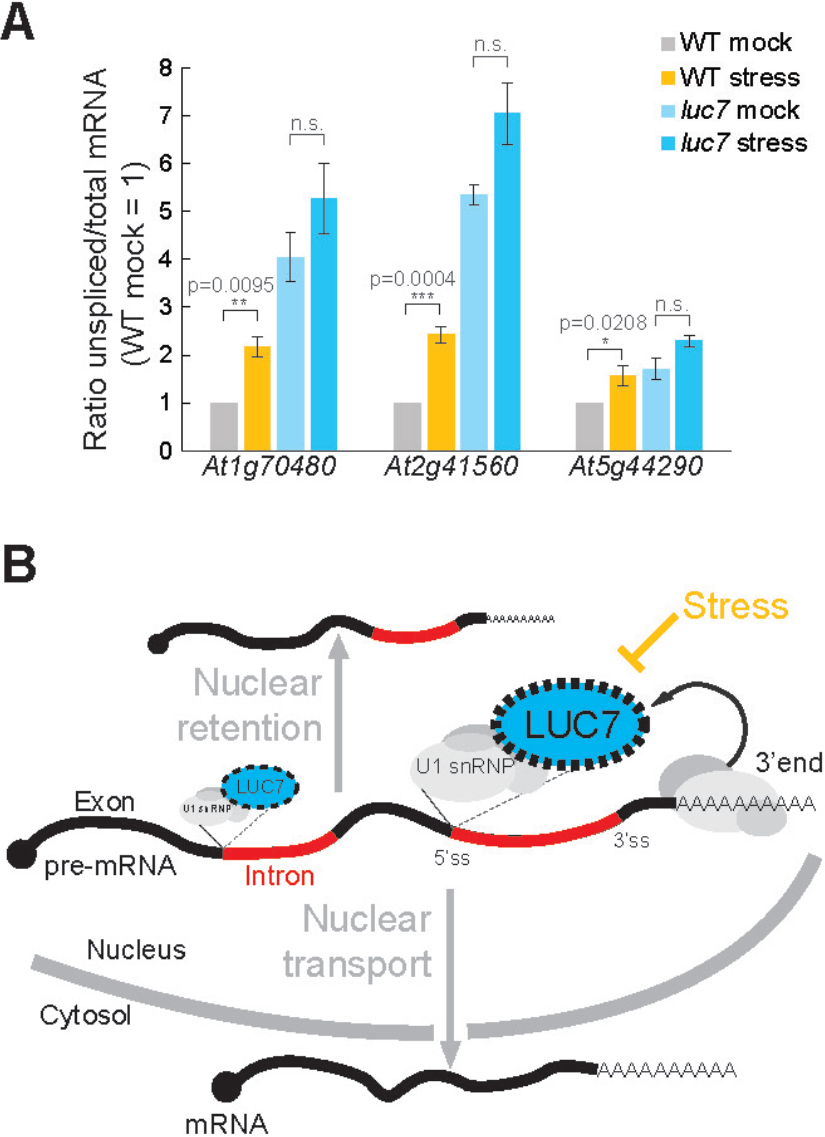
Splicing of LUC7 dependent introns can be modulated by stress. **A:** Seven days old WT and *luc7* triple mutant seedlings were exposed to cold for 60 min. Splicing ratios (unspliced/total RNA) of four genes featuring a LUC7-dependent intron was analyzed by qPCR. A T-test was performed for statistical analysis. **B:** Model for the proposed function of LUC7 in Arabidopsis.

## Discussion

### Functions of the Arabidopsis U1 snRNP component LUC7

For this study, we generated an Arabidopsis triple mutant deficient in the U1 snRNP components LUC7 and dissected the genome-wide effects of LUC7s impairments on the Arabidopsis transcriptome. Our results show that LUC7 proteins are *bona-fide* U1 components acting mainly redundantly. The reduction of U1 function in the *luc7* triple mutant affects constitutive splicing. A large number of introns are retained in *luc7* triple mutant, suggesting that without a proper recognition of the 5’ss, splicing of the affected introns is impaired. Our results also show that exon-skipping events are impaired in *luc7* triple mutant, revealing that a functional plant U1 snRNP is essential for exon definition. In addition, we show that *luc7* triple mutant affect alternative splicing also by influencing events of alternative 5’ and 3’ splice site. This implies that the U1 snRNP does not only affect 5’ splice site usage, it might also indirectly regulate usage of 3’ splice sites via its interaction with U2AFs and the U2 snRNP (Hoffman and Grabowski, 1992; Shao et al., 2012). The functions of LUC7 proteins on the Arabidopsis transcriptome are likely to be underestimated, because misspliced mRNAs in *luc7* mutants might contain hallmarks of NMD and are therefore rapidly turned over and escape detection. Analysis of *luc7* mutants combined with mutations in NMD factors would help to uncover the full set of splicing events affected by LUC7. Furthermore, we found that in our RNA-seq experiments that while the chosen *luc7rl* allele is a RNA-null allele, the *luc7a* and *luc7b* alleles still produced mRNAs that might be translated into truncated proteins. Hence, we can not exclude that a true *luc7* null mutant might exhibit even more severe mutant phenotypes and splicing defects. One has also to consider that U1 snRNP independent splicing has been described in animals, indicating that not all introns require the U1 complex for efficient intron removal (Fukumura et al., 2009). The degree of U1-independent splicing in plants remains to be elucidated.

Duplications among genes encoding for U1 snRNP proteins, such as the LUC7 genes, open up a possibility for sub- and neofunctionalization of U1 accessory proteins. Furthermore, the Arabidopsis genome encodes 14 potential U1 snRNAs, which slightly differ in sequence (Wang and Brendel, 2004). Therefore, the plant U1 snRNP presumably does not exist as a single complex, but might exist as different sub-complexes exhibiting distinct specificities and functions. In metazoans, the existence of at least four different U1 snRNP subcomplex has been suggested (Hernandez et al., 2009; Guiro and O’Reilly, 2015). Specific combinations of plant U1 protein family members and U1 snRNAs could generate an even higher number of such U1 subcomplexes, which could be responsible for specific splicing events. Our results show that LUC7 can act redundantly, but can also fulfill specific functions, suggesting that LUC7 complexes specifically act on certain pre-mRNAs. In this regard, it is important to note that an additional short stretch of amino acids separates the two zinc-finger domains in LUCA and LUC7B (Figure S1). Changing the space in between RNA binding domains affects substrate specificities and could explain different specificities among LUC7 proteins (Chen and Varani, 2013).

### LUC7 function in terminal intron splicing

Interestingly, *luc7* triple mutant showed a significant higher retention rate of terminal introns compared to first or middle introns. This was surprising because LUC7 was initially found to act in concert with the CBC, a complex involved in the removal of cap-proximal first introns, but not of last introns (Lewis et al., 1996). We found LUC7 in a complex with the CBC and the CBC-associated protein SE also in Arabidopsis. However, the functional significance of the of LUC7-CBC/SE interaction remains to be established.

Often, the removal of terminal introns is coupled to polyadenylation (Cooke et al., 1999; Cooke and Alwine, 2002; Rigo and Martinson, 2008). Interestingly, we detected components involved in RNA 3’end processing or polyA-binding as part of LUC7A complexes, suggesting that such interactions may contribute to the specific functions of LUC7 in terminal intron splicing. We found LUC7 in complexes with RBP47C, a polyA-binding protein of unknown function (Lorkovic et al., 2000), and LARP6, which is targeted to 3’ends of mRNA through interaction with polyA-binding protein 2 (PAB2) (Merret et al., 2013). Interestingly, we also found an SPT6-like transcription factor associated with LUC7 complexes. SPT6 binds the pol II C-terminal domain (CTD) phosphorylated at serine 2 (Ser2P), which accumulates at 3’end of genes (Kaplan et al., 2000; Sun et al., 2010). None of these LUC7 complex components has been studied functionally and it will be a major effort for future studies to determine the function of these proteins in terminal intron splicing.

### Possible functions of regulated intron retention for plant stress responses

We found that splicing of LUC7-dependent introns is required for transport of mRNAs from the nucleus to the cytosol. The fact that we cannot detect unspliced transcript in the cytosol suggests a nuclear retention mechanism for such mRNAs. One possibility is that LUC7-dependent introns might contain binding sites for specific trans-regulatory factors that upon binding inhibit export. Polypyrimidine tract-binding protein 1 (PTB1) is a candidate for such a trans-regulatory protein, because binding of PTB1 to introns represses nuclear export of certain RNAs (Yap et al., 2012; Roy et al., 2013).

Nuclear retention of unspliced mRNAs might be a much more general mechanism to escape NMD and to regulate gene expression (Marquez et al., 2015; Wong et al., 2016). In plants, some specific transcript isoforms have been detected only in the nucleus, but not in the cytosol (Gohring et al., 2014). Also in metazoans, intron retention might have a more general role in regulating gene expression (Yap et al., 2012; Braunschweig et al., 2014; Pimentel et al., 2016; Naro et al., 2017). The so-called detained introns are evolutionary conserved, NMD insensitive and retained in the nucleus (Boutz et al., 2015). The functional importance of intron retention was also suggested in the fern *Marsilea vestita*, in which many mRNAs contain introns that are only spliced shortly before gametophyte development (Boothby et al., 2013).

We found that splicing of LUC7 dependent introns can be modulated by cold stress. Because retention of these introns causes nuclear trapping, it is prompting to speculate that environmental cues affect splicing and nuclear retention of mRNAs. Such a mechanism would regulate the amount of translatable mRNAs in the cytosol in a cost-efficient and rapid manner (Figure 8B). Since the stress-dependent regulation of splicing of LUC7-dependent introns is lost in *luc7* mutants, one can expect that LUC7 function might be regulated under stress conditions. Interestingly, the RS domains of LUC7 proteins are phosphorylated and we identified three kinases as potential LUC7A interactors (Heazlewood et al., 2008; Durek et al., 2010). In addition, stress signaling triggered by the phytohormone ABA causes differential phosphorylation of several splicing factors (Umezawa et al., 2013; Wang et al., 2013). Whether stress-induced changes in phosphorylation play a role in regulating LUC7 proteins and whether the LUC7-interacting kinases here identified are involved in this process remains to be elucidated.

## Material and Methods

### Plant material and growth conditions

All mutants were in the Columbia-0 (Col-0) background. *luc7a-1* (SAIL_596_H02) and *luc7a-2* (SAIL_776_F02), *luc7b-1* (SALK_144681), *luc7rl-1* (SALK_077718) and *luc7rl-2* (SALK_130892C) were isolated by PCR-based genotyping (Table S7). *luc7* double and triple mutants were generate by crossing individual mutants. All other mutants used in this study (*abh1-285*, *cbp20-1*, *se*-1, *lba-1* and *upf3-1*) were described elsewhere (Prigge and Wagner, 2001; Papp et al., 2004; Hori and Watanabe, 2005; Yoine et al., 2006; Laubinger et al., 2008). The line expressing GFP was generated using the vector pBinarGFP and was kindly provided by Dr. Andreas Wachter (Wachter et al., 2007). For complementation analyses, *pLUC7A:LUC7A-FLAG*, *pLUC7B:LUC7B-FLAG*, *pLUC7RL:LUC7RL-FLAG* and *pLUC7A:LUC7A-YFP* constructs were introduced in *luc7* triple mutant by Agrobacterium-mediated transformation (Clough and Bent, 1998). All plants were grown on soil in long days conditions (16-h light/8-h dark) at 20°C/18°C day/night. The size of *luc7* mutants was assessed by measuring the longest rosette leaf after 21 days. For all molecular studies, seeds were surface-sterilized, plated on 1/2 MS medium with 0.8% phytoagar and grown for 7 days in continuous light at 22°C. For the cold treatment, plates with *Arabidopsis* seedlings were transferred to ice-water for 60 min. For the root growth assay, 4 days old seedlings growing on vertical plates were transferred to mock plates or plates containing indicated amount of NaCl and grown for more 11 days vertically. Root growth rate per day was assessed by measuring in ImageJ the root length in the days 2 and 9 after transfer. For ABA sensitivity assays, seedlings were grown for 10 days on 1/2 MS plates supplemented with 0.8 % phytoagar, 1 % sucrose and indicated amounts of ABA (+) (Sigma - A4906). For cold stress experiments, seeds were grown at 20°C for 5 days and then transferred to 8°C for two weeks.

### Plasmid constructions and transient expression analyses

For the expression of C-terminal FLAG- and YFP-tagged LUC7 proteins expressed from their endogenous regulatory elements, 2100 bp, 4120 bp and 2106 bp upstream of the ATG start codon of *LUC7A*, *LUC7B* and *LUC7RL*, respectively, to the last coding nucleotide were PCR-amplified and subcloned in pCR8/GW/TOPO® (Invitrogen). Oligonucleotides are listed in Table S7. Entry clones were recombined with pGWB10 and pGWB540 using Gateway LR clonase II (Invitrogen) to generate binary plasmids containing *pLUC7A:LUC7A-FLAG*, *pLUC7B:LUC7B-FLAG*, *pLUC7RL:LUC7RL-FLAG* and *pLUC7A:LUC7A-YFP*. For the co-localization studies, entry vector containing the coding sequence of U1-70k was recombined with pGWB654 for the expression of *p35S:U1-70k-mRFP* (Nakagawa et al., 2007). Agrobacterium-mediated transient transformation of *Nicotiana benthamiana* plants was conducted as following. Overnight Agrobacterium culture were diluted in 1:6 and grown for 4 hours at 28°C. After centrifugation, pellets were resuspended in infiltration medium (10 mM MgCl_2_, 10 mM MES-KOH pH 5.6, 100 µM Acetosyringone). The OD 600nm was adjusted to 0.6-0.8 and samples were mixed when required. *N. benthamiana* were infiltrated and subcellular localization was checked after 3 days. Subcellular localization of fluorescent proteins was analyzed by confocal microscopy using a Leica TCS SP8.

### Phylogenetic analysis

AthLUC7A (AT3G03340) protein sequence was analyzed in Interpro (https://www.ebi.ac.uk/interpro/) to retrieve the Interpro ID for the conserved Luc7-related domain (IPR004882). The sequence for *Saccharomyces cerevisiae* (strain ATCC 204508_S288c) was obtained in Interpro. Plants sequences were extracted using BioMart selecting for the protein domain IPR004882 on Ensembl Plants (http://plants.ensembl.org/). The following genomes were included in our analyses: *Amborella trichopoda* (AMTR1.0 (2014-01-AGD)); *Arabidopsis thaliana* (TAIR10 (2010-09-TAIR10)); *Brachypodium distachyon* (v1.0); *Chlamydomonas reinhardtii* (v3.1 (2007-11-ENA)); *Physcomitrella patens* (ASM242v1 (2011-03-Phypa1.6)); *Selaginella moellendorffii* (v1.0 (2011-05-ENA)); *Oryza sativa Japonica* (IRGSP-1.0); and *Ostreococcus lucimarinus* genes (ASM9206v1). The phylogenetic analysis was performed in Seaview (Version 4.6.1) using Muscle for sequence alignment. Maximum likehood (PhYML) was employed with 1000 bootstraps (Gouy et al., 2010).

### RNA extractions, RT-PCR and qRT-PCR

RNAs extractions were performed with Direct-zol™ RNA MiniPrep Kit (Zymo Research). Total RNAs were treated with DNAse I and cDNA synthesis carried out with RevertAid First Strand cDNA Synthesis Kit (Thermo Scientific) using usually oligo dT primers or a mixture of hexamer and miRNA-specific stem-loop primers (Table S7). Standard PCRs for the splicing analysis were performed with DreamTaq DNA Polymerase (Thermo Scientific). Quantitative RT-PCR (qRT-PCR) was performed using the Maxima SYBR Green (Thermo Scientific) in a Bio-Rad CFX 384. For all qPCR-primers, primer efficiencies were determined by a serial dilution of cDNA template. The relative expressions were calculated using the 2^(-ΔΔCT) method with *PP2A* or *ACTIN* as control. For the qRT-PCR to detect splicing ratio changes under cold condition, the ratio 2^(-ΔCT_unspliced_)/2^(-ΔCT_total RNA_) was calculated separately for each replicate and t-test was performed before calculating the relative to WT mock. Oligonucleotides are listed in Table S7. For RNA-sequencing analysis, polyA RNAs were enriched from 4 µg of total RNAs using NEBNext Oligo d(T)_25_ Magnetic Beads (New England Biolabs). The libraries were prepared using ScriptSeq™ Plant Leaf kit (Epicentre) following the manufacturer’s instruction. Single end sequencing was performed on an Illumina HiSeq2000. Sequencing data were deposited at Gene Expression Omnibus under accession number GSE98779.

### RNA-seq libraries: Mapping, differential expression analysis and splicing analysis

RNA-seq reads for each replicate were aligned against the *Arabidopsis thaliana* reference sequence (TAIR10) using tophat (v2.0.10, -p2, -a 10, -g 10, -N 10, --read-edit-dist 10, --library-type fr-secondstrand, --segment-length 31, -G TAIR10.gff). Next, cufflinks (version 2.2.1) was used to extract FPKM counts for each expressed transcript generating a new annotation file (transcripts.gtf), where the coordinates of each expressed transcript can be found. Cuffcompare (version 2.2.1) was then used to generate a non-redundant annotation file containing all reference transcripts in addition to new transcripts expressed in at least one of the nine samples (cuffcmp.combined.gtf). The differential expression analysis was performed with cuffdiff (version 2.2.1) between wt/*luc7* triple using the annotation file generated by cuffcompare (false discovery rate (FDR) < 0.05 and fold change (FC) > 2). For the splicing analysis, the same alignment files generated by tophat and annotation files generated by cuffcompare (cuffcmp.combined.gtf) were used as input for MATS (version 3.0.8) in order to test for differentially spliced transcripts (Shen et al., 2014).

### Subcellular fractionation

Two grams of seedlings were ground in N_2_ liquid and resuspended in 4 ml of Honda buffer (0.44 M sucrose, 1.25% Ficoll 400, 2.5% Dextran T40, 20 mM HEPES KOH pH 7.4, 10 mM MgCl_2_, 0.5% Triton X-100, 5 mM DTT, 1 mM PMSF, protease inhibitor cocktail (Roche) supplemented with 40U/ml of Ribolock®). The homogenate was filtered through 2 layer of Miracloth, which was washed with 1ml of Honda Buffer. From the filtrate, 300 µl was removed as “total” fraction and kept on ice. Filtrates were centrifuged at 1,500 *g* for 10 min, 4°C for pelleting nuclei and supernatants were transferred to a new tube. Supernatants were centrifuged at 13 000 x g, 4 °C, 15 min and 300 µl were kept on ice as cytoplasmic fraction. Nuclei pellets were washed five times in 1 ml of Honda buffer (supplemented with 8U/ml of Ribolock®, centrifugation at 1,800 *g* for 5 min. The final pellet was resuspended in 300 µl of Honda buffer. To all the fractions (total, cytoplasmic and nuclei), 900 µl of TRI Reagent (Sigma) was added. After homogenization, 180 µl of chloroform was added and samples were incubated at room temperature for 10 min. After centrifugation at 10 0000 rpm for 20 min, 4°C, the aqueous phase were transferred to a new tube and RNA extracted with Direct-zol™ RNA MiniPrep Kit (Zymo Research). The organic phase was collected and proteins were isolated according to manufacturer’s instructions. cDNA synthesis with random primes was performed as above. Proteins extracted were analyzed by standard western blot techniques using the following antibodies: H3 (∼ 17 KDa / ab 1791, Abcam) and 60S ribosomal (∼ 23,7-29 KDa / L13, Agrisera).

### RNA immunoprecipitation

RNA immunoprecipitation (RIP) using WT and a *pLUC7A:LUC7A-eYFP* rescue line was performed as described elsewhere with minor modifications (Rowley et al., 2013; Xing et al., 2015). Isolated nuclei were sonicated in nuclear lysis buffer in a Covaris E220 (Duty Cycle: 20%; Peak intensity: 140; Cycles per Burst: 200; Cycle time: 3’). RNAs were extracted using RNeasy Plant Mini Kit (QIAGEN) following the manufacturer’s instructions. The RNA were treated with DNAseI (Thermo Scientific) and samples were split in half for the (-)RT reaction. cDNA synthesis were perform with SuperScript™ III Reverse Transcriptase (Invitrogen). qRT-PCRs were performed with QuantiNova™ SYBRR Green PCR (QIAGEN).

### GO Analysis

GO analysis was performed in Bar Utoronto (http://bar.utoronto.ca/ntools/cgi-bin/ntools_classification_superviewer.cgi).

### Protein complex purification and mass spectrometry (MS) Analysis

LUC7A immunoprecipitation was performed using a complemented line *pLUC7A:LUC7A-eYFP* (line 20.3.1) and a transgenic *p35S:GFP* and WT as controls. Four independent biological replicates were performed. Seedlings (4 g) were ground in N_2_ liquid and respuspended in 1 volumes of extraction buffer (50 mM Tris-Cl pH 7.5, 100 mM NaCl, 0.5 % Triton X-100, 5 % Glycerol, 1 mM PMSF, 100 µM MG132, Complete Protease Inhibitor Cocktail EDTA-free [Roche] and Plant specific protease Inhibitor, Sigma P9599). After thawing, samples were incubated on ice for 30 min, centrifuged at 3220 rcf for 30 min at 4°C and filtrated with two layers of Miracloth. For each immunoprecipitation, 20 µl of GFP-trap (Chromotek) was washed twice with 1 ml of washing buffer (50 mM Tris-Cl pH 7.5, 100 mM NaCl, 0.2 % Triton X-100) and once with 0.5 ml of IP buffer. For each replicate, the same amount of plant extracts (∼5 ml) were incubated with GFP-trap and incubated on a rotating wheel at 4°C for 3 hours. Samples were centrifuged at 800-2000 rcf for 1-2 min and the supernatant discarded. GFP-beads were resuspended in 1 ml of washing buffer, transferred into a new tube and washed 4 to 5 times. Then, beads were ressuspended in ∼40 µl of 2x Laemmli Buffer and incubated at 80°C for 10 min. Short gel purifications (SDS-PAGE) were performed and gels slices were digested with Trypsin. LC-MS/MS analyses were performed in two mass spectrometer. Samples from R10 to R14 were analyzed on a Proxeon Easy-nLC coupled to Orbitrap Elite (method: 90min, Top10, HCD). Samples from R15 to R17 were analysed on a Proxeon Easy-nLC coupled to OrbitrapXL (method: 90min, Top10, CID). Samples from R18 to R20 analysis on a Proxeon Easy-nLC coupled to OrbitrapXL (method: 130min, Top10, CID). All the replicates were processed together on MaxQuant software (Version 1.5.2.8. with integrated Andromeda Peptide search engine) with a setting of 1% FDR and the spectra were searched against an *Arabidopsis thaliana* Uniprot database (UP000006548_3702_complete_20151023.fasta). All peptides identified are listed in Supplementary Table S1 and raw data were deposited publically (accession PXD006127). For co-immunoprecipitation experiments shown in Figure S4, experiments were conducted as described above and IPed protein fractions were analyzed using SDS-PAGE followed by detection with GFP-(Roche), SE-(Agrisera) and CBP80-(Agrisera) specific antibodies.

## Acknowledgments

This work was supported by the Deutsche Forschungsgemeinschaft DFG (to S.L., LA2633-4/1), Coordenação de Aperfeiçoamento de Pessoal de Nível Superior (CAPES - Brazil) for doctoral fellowship (to M.F.A.), the Max Planck Society (to K.S.), the Max Planck Society Chemical Genomics Centre (CGC) through its supporting companies AstraZeneca, Bayer CropScience, Bayer Healthcare, Boehringer-Ingelheim and Merck (to S.L). We are grateful to Andreas Wachter (ZMBP, University of Tuebingen, Germany) and members of the lab for critical reading of the manuscript, Christa Lanz for her invaluable help with Illumina sequencing, and Johanna Schröter and her team for excellent care of our plants, Anja Hoffmann for excellent technical assistance, and Andreas Wachter (ZMBP, University of Tuebingen, Germany), Tsuyoshi Nakagawa (Department of Molecular and Functional Genomics, Center for Integrated Research in Science, Shimane University, Matsue, Japan) and the Notthingham Arabidopsis Stock Centre for providing seeds and DNA constructs.

## Author contributions

M.d.F.A. and S.L. designed research. M.d.F.A., A.G.F.-M., I.D-B. and S.L. performed experiments. E.-M. W., M.d.F.A., A.G.F.-M., I.D-B., B.M., K.S. and S.L. analyzed the data. M.d.F.A. and S.L. wrote the manuscript with contributions from all authors.

## Conflict of interest

The authors declare no conflict of interest.

## Supplementary Material

Figure S1-S8, Table S1-S7

