## Supplementary Materials for "The U1 snRNP subunit LUC7 controls plant development and stress response through alternative splicing regulation"

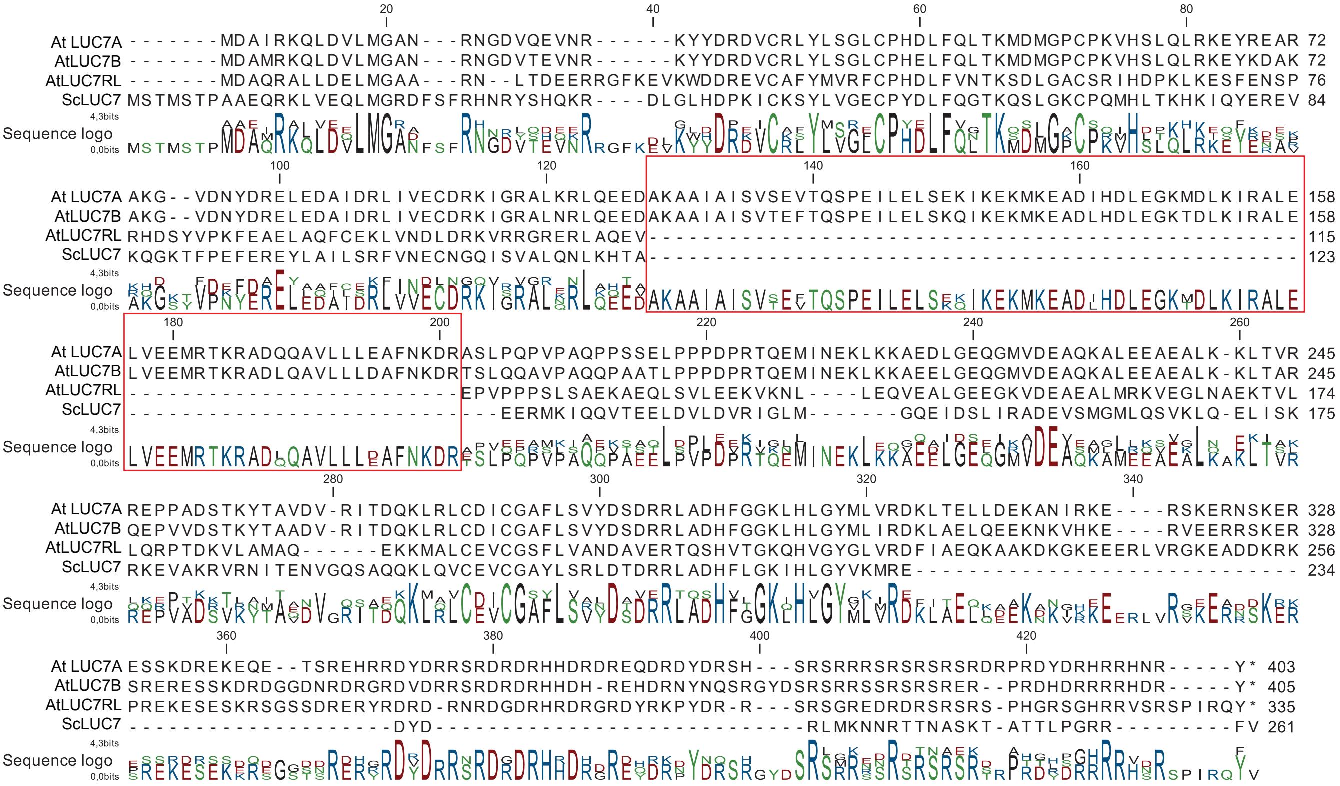


**FIGURE S1**

Figure S1: Protein alignment *A. thaliana* LUC7 proteins and *S. cerevisiae* LUC7. The red box indicates a stretch of amino acids specific for LUC7A and LUC7B.

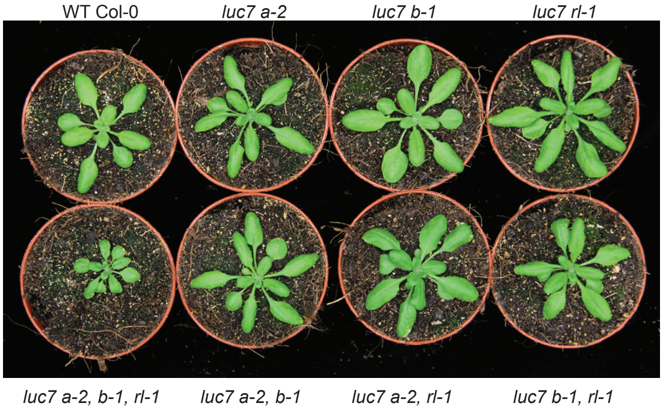


**FIGURE S2**

Figure S2: Growth phenotypes of WT, *luc7* single, double and triple mutants.

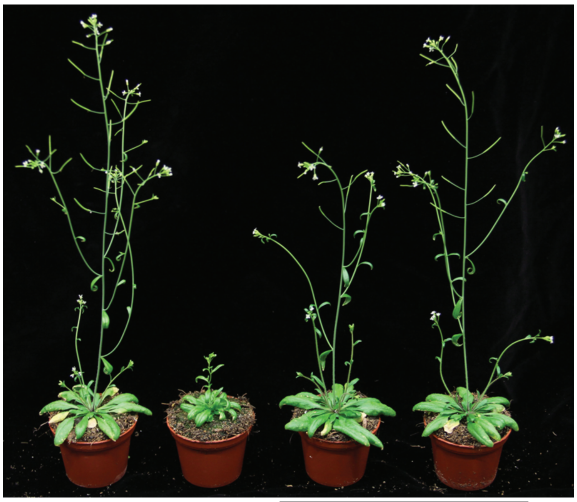


**FIGURE S3**

Figure S3: Growth phenotypes of WT, *luc7* triple mutants and *luc7* triple mutants containing a *LUC7A:LUC7A-GFP* rescue construct.

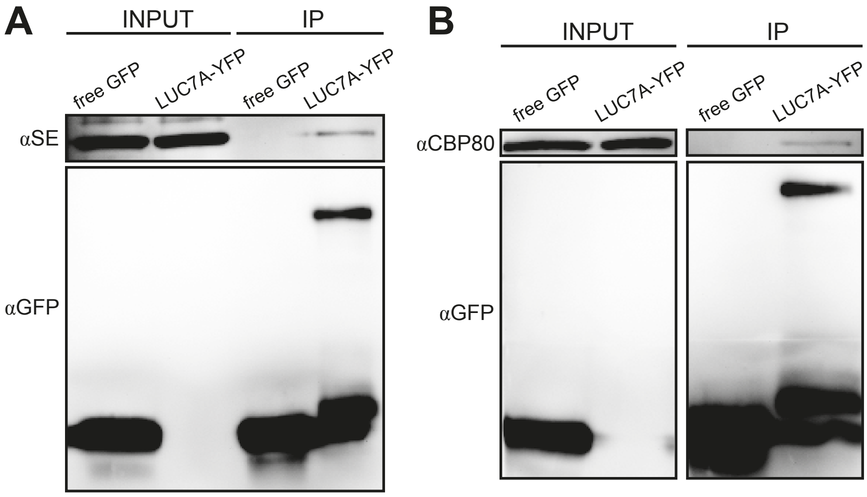


**FIGURE S4**

Figure S4: Interaction test between LUC7A and SE (A) and CBP80 (B). LUC7A-YFP was affinity purified from transgenic plants and co-immunoprecipitated were analyzed by protein blot with GFP-, SE- and CBP80-secific antibodies. Please note that the LUC7A-YFP fusion protein is detectable only in IP fractions.

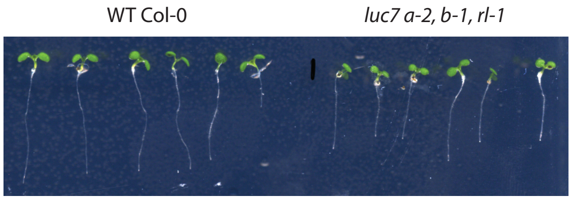


**FIGURE S5**

Figure S5: Growth phenotypes of seven-day old WT and *luc7* triple mutant seedlings grown on MS media.

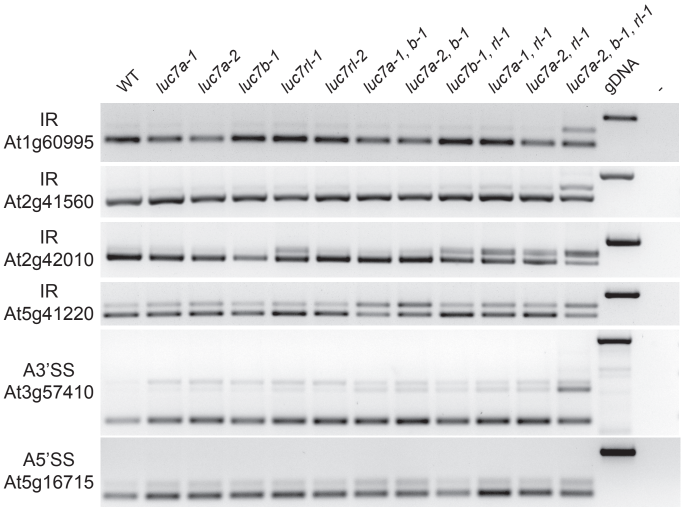


**FIGURE S6**

Figure S6: RT-PCR splicing analysis in WT, *luc7* single, double and triple mutants.

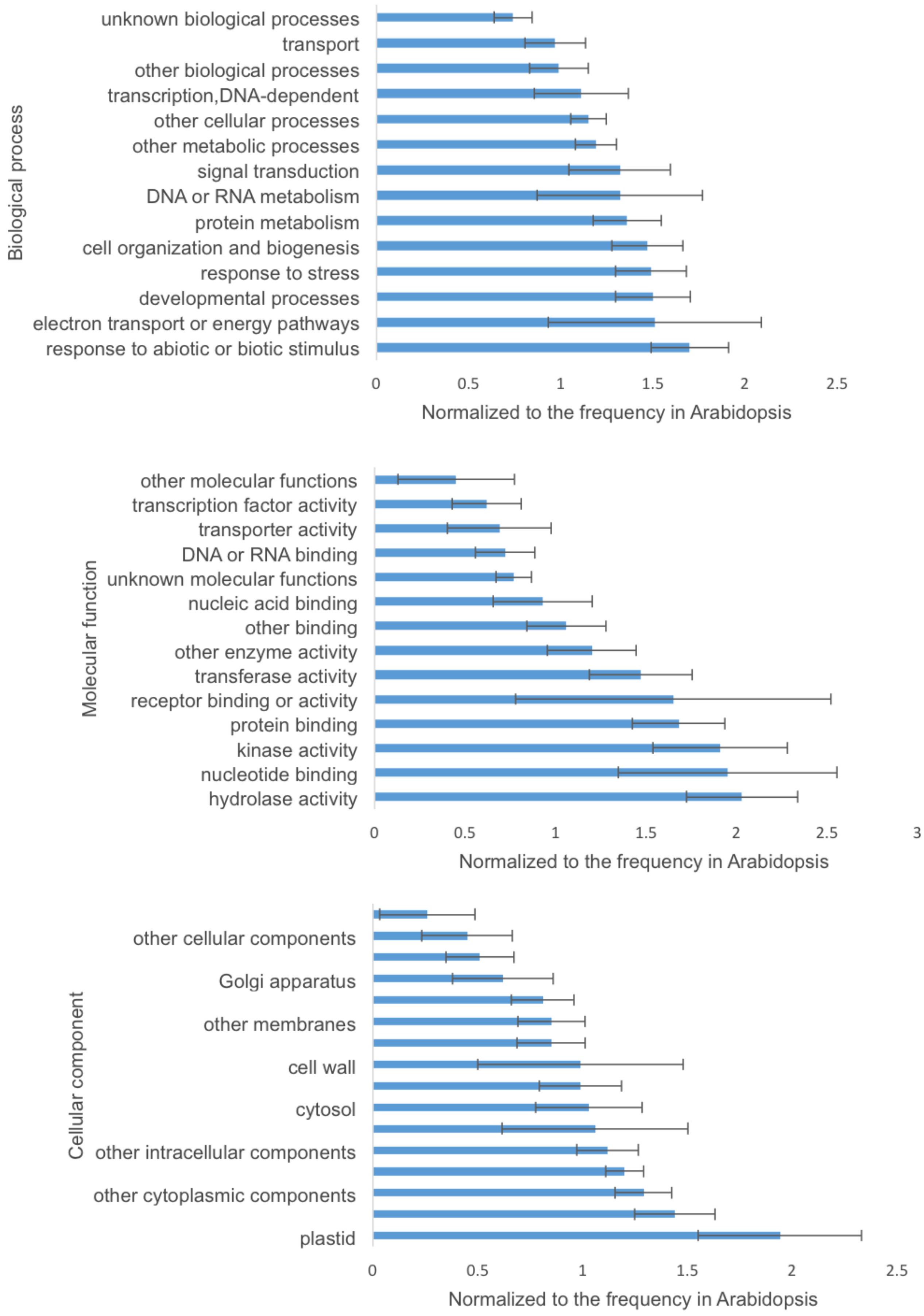


**FIGURE S7**

Figure S7: GO analysis of genes, which mRNAs contain retained introns in *luc7* triple mutants.

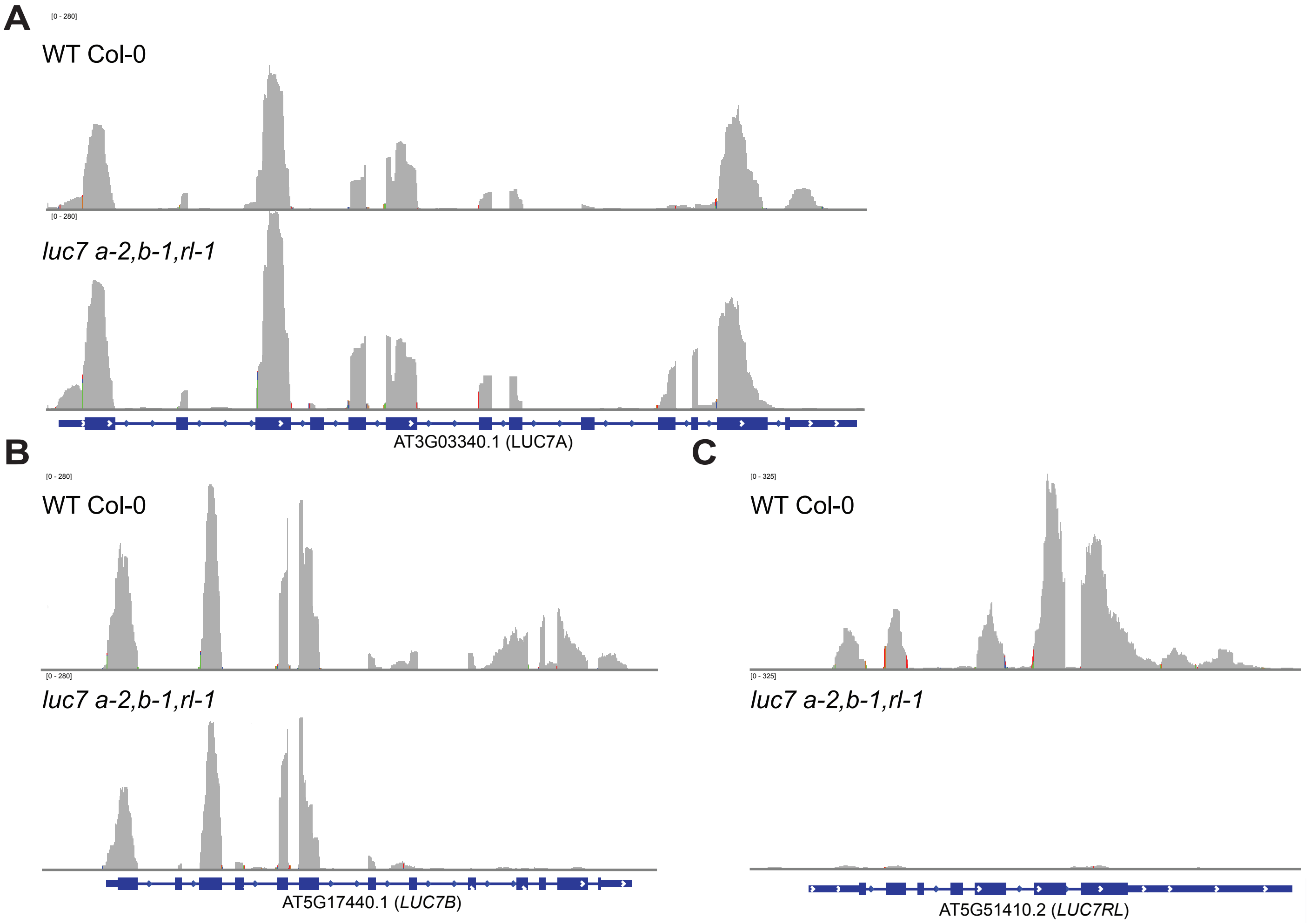


### FIGURE S8

Figure S8: Coverage plots of RNA-seq reads at the *LUC7A* (A), *LUC7B* (B) and *LUC7RL* (C) locus in WT and *luc7* triple mutants.
